# Structural basis of αE-catenin–F-actin catch bond behavior

**DOI:** 10.1101/2020.07.13.201236

**Authors:** Xiao-Ping Xu, Sabine Pokutta, Miguel Torres, Mark F. Swift, Dorit Hanein, Niels Volkmann, William I. Weis

## Abstract

Cell-cell and cell-matrix junctions transmit mechanical forces during tissue morphogenesis and homeostasis. α-Catenin links cell-cell adhesion complexes to the actin cytoskeleton, and mechanical load strengthens its binding to F-actin in a direction-sensitive manner. This so-called catch bond behavior is described by a model in which force promotes a transition between weak and strong actin-bound states. We describe the cryo-electron microscopy structure of the F-actin-bound αE-catenin actin-binding domain, which in solution forms a 5-helix bundle. Upon binding to actin, the first helix of the bundle dissociates and the remaining four helices and connecting loops rearrange to form the interface with actin. Deletion of the N-terminal helix produces strong actin binding in the absence of force. Our analysis explains how mechanical force applied to αE-catenin or its homolog vinculin favors the strongly bound state, and the dependence of catch bond strength on the direction of applied force.

## INTRODUCTION

The development and maintenance of multicellular organisms depends upon specific adhesion between cells. Morphogenetic movements of sheets of cells are driven by changes in the cytoskeleton of individual cells that are linked to adjacent cells by adhesion molecules. Tissue integrity depends upon response of these adhesive structures to external mechanical perturbation (Guillot and Lecuit, 2013; Ladoux and Mege, 2017). Cell-cell junctions, including the adherens junctions (AJ), transmit mechanical forces between cells. In AJs, the extracellular domains of cadherins mediate homophilic cell-cell contact, and their cytoplasmic domains are linked to the actin cytoskeleton by β-catenin and α-catenin (Meng and Takeichi, 2009; Shapiro and Weis, 2009). Specifically, β-catenin binds to the cytoplasmic tails of cadherins and to α-catenin; α-catenin binds to β-catenin and to filamentous (F-)actin (Figure 1a). This architecture enables forces generated by actomyosin constriction to be transmitted to neighboring cells during morphogenesis, and conversely allows the actin cytoskeleton to respond to external loads. Similarly, in cell-extracellular matrix adhesions, the extracellular domains of integrins bind to components of the extracellular matrix. The cytoplasmic protein talin binds to integrins and to vinculin, a homolog of α-catenin.

**Figure 1.**
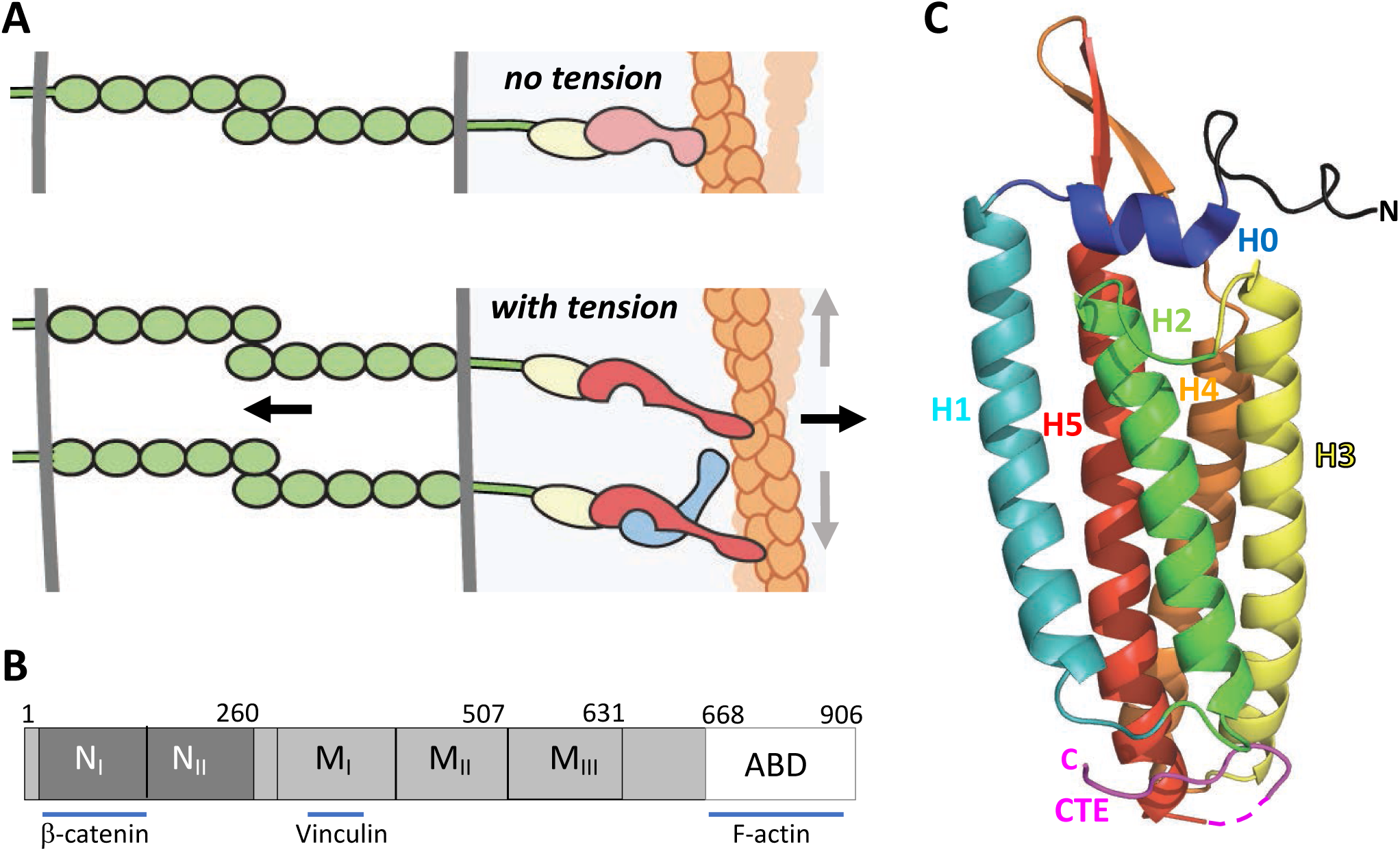
α-Catenin in adherens junctions. **A**. Schematic of AJ and the role of αE-catenin in the connection to actin cytoskeleton. The extracellular region of cadherins (green) bind to one another between cells, and their cytoplasmic domains bind to β-catenin (yellow). β-Catenin binds to α-catenin (pink/red), which binds to F-actin (orange) weakly in the absence of force (top panel). Tension (indicated by arrows) favors the strong actin-binding state of α-catenin, and also produces conformational changes that lead to recruitment of vinculin (light blue). While the net direction of the force is likely perpendicular to the junction (black arrows), there will be local force components along the mixed-polarity filaments towards their pointed (-) ends through actomyosin contractility (grey arrows). **B**. Primary structure of αE-catenin. **C**. Crystal structure of αE-catenin ABD (Ishiyama et al., 2018); the five helices H1-H5, the N-terminal capping helix H0, and the C-terminal extension (CTE) are labeled.

α-Catenin appears to be the major sensor of mechanical force in the AJ. Tension on cadherins depends upon actomyosin activity, β-catenin and α-catenin (Borghi et al., 2012). As AJs develop, tension placed on α-catenin promotes conformational changes that enable it to bind to its paralog vinculin (Barrick et al., 2018; Kim et al., 2016; le Duc et al., 2010; Li et al., 2015; Maki et al., 2016; Maki et al., 2018; Terekhova et al., 2019), whose actin-binding activity further strengthens the cytoskeletal linkage (Thomas et al., 2013). In the mature AJ, actin bundles of mixed polarity run parallel to the junction (Hirokawa et al., 1983) and lie in close apposition to the membrane (Buckley et al., 2014). Single molecule force measurements of the cadherin–catenin complex binding to actin that employed an optical trap setup revealed that the complex displays catch bond behavior, wherein the interaction with actin is strengthened under mechanical load (Buckley et al., 2014). This property was observed subsequently in α-catenin and full-length vinculin themselves, as well as the actin-binding domain (ABD) of vinculin, indicating that the homologous ABDs of these proteins confer catch bond behavior (Abore et al., 2020; Huang et al., 2017). In both proteins, the catch bond is asymmetric: force directed towards the (-) end of the actin filament results in a longer lived bond than when force is directed towards the (+) end (Abore et al., 2020; Huang et al., 2017).

α-Catenin has three major domains: an N-terminal β-catenin binding domain, a middle (M) domain, followed by a flexible linker to the C-terminal ABD (Figure 1b). The three-dimensional structures of the ABD from αE(epithelial)- and αN(neuronal)-catenins have been determined (Ishiyama et al., 2018; Ishiyama et al., 2013), and the ABD has also been visualized in the crystal structure of a nearly full-length αE-catenin (Rangarajan and Izard, 2013). These structures reveal that the ABD comprises a bundle of five helices, preceded by a short N-terminal helix (designated H0) that sits on top of the bundle (Figure 1c). Helices 2, 3, 4 and 5 (H2-5) form an antiparallel four-helix bundle in which hydrophobic residues from each helix contribute to a hydrophobic core. Helix 1 interacts with the side of the four helix H2-H5 bundle. A long C-terminal extension (CTE), residues 844-906, follows H5. In different structures the CTE adopts different conformations and is partly disordered. The vinculin ABD likewise adopts a similar 5 helix bundle architecture, albeit with a shorter H1 and no H0 (Bakolitsa et al., 2004; Bakolitsa et al., 1999; Borgon et al., 2004).

In the optical trap data, the distribution of bound lifetimes of the cadherin/catenin complex or vinculin at any particular force follows a bi-exponential distribution, indicating that there are two distinct actin-bound states, weak and strong (Buckley et al., 2014; Huang et al., 2017). The population of longer lifetimes increases with force, and modeling of these data indicated that the catch bond behavior arises because force enhances interconversion of the weakly- to the strongly-bound state (Buckley et al., 2014). These observations explain why binding of the cadherin–catenin complex to actin in solution, *i*.*e*., under no external load, is weak; force shifts the equilibrium between the weakly and strongly bound states and thereby produces tighter binding (Buckley et al., 2014; Yamada et al., 2005).

While the catch bonding of αE-catenin and vinculin to actin is established, to date there has been no molecular explanation of how force changes the structure of their ABDs to promote strong binding. Recent work in solution demonstrated removal of H0 from the ABD of αE-catenin enhances its affinity for F-actin, suggesting that force-dependent removal of this structural element is important for catch bonding (Ishiyama et al., 2018). However, vinculin lacks H0 yet also displays catch bond behavior (Bakolitsa et al., 2004; Bakolitsa et al., 1999; Borgon et al., 2004; Huang et al., 2017). Here, we present the structure of the αE-catenin ABD lacking H0 bound to F-actin obtained by cryo-electron microscopy (cryo-EM). Comparison of the crystal structure and the actin-bound form of the complete αE-catenin ABD, as well as the structures of the vinculin ABD free or bound to actin (Bakolitsa et al., 2004; Bakolitsa et al., 1999; Borgon et al., 2004; Mei et al., 2020), provides an explanation for the weak to strong actin-binding transition, and biochemical and mutational data support this model.

## RESULTS

### Structure of αE-catenin ABD bound to F-actin

To understand the αE-catenin-actin filament interaction in molecular detail, we obtained a three-dimensional cryo-EM reconstruction of actin filaments bound to a truncated αE-catenin ABD (residues 671-906) at 3.8 Å resolution (Fig. 2, Figure 2-figure supplement 1), Table 1). This construct deletes the first half of, and thereby destabilizes, the short H0, and binds 4.5x more strongly than the complete ABD (residues 666-906) (Table 2, Table 2-figure supplement 1) (Ishiyama et al., 2018). Consistent with the previously described cooperative binding by this construct (Hansen et al., 2013), we observed either bare actin filaments or stretches of filaments continuously bound by αE-ABD. Using the same 10 μM concentration, we were unable to observe binding of the complete ABD to actin filaments in the electron microscope. Given the K_D_ values of these two constructs, small changes in concentration likely have a large effect on actin decoration. Indeed, a comparable structure using an ABD construct spanning residues 664-906 was produced using a higher concentration of ABD (Mei et al., 2020).

**Table 1.** Cryo-EM data collection and model statistics.

|  |  |
| --- | --- |
| **Data collection** |  |
| Microscope | Titan Krios |
| Voltage [kV] | 300 |
| Detector | Falcon II |
| Magnification | 75,000 |
| Total exposure [e<sup>-</sup>/Å<sup>2</sup>] | 60 |
| Pixel size [Å] | 1.035 |
| Defocus range [μm] | -0.8 to -2.8 |
| **Data processing** |  |
| Images used | 4769 |
| Initial segments | 728,331 |
| Final segments | 422,822 |
| *Helical symmetry* |  |
| Rise [Å] | 27.4 |
| Twist [°] | -166.9 |
| Resolution [Å] | 3.56 |
| FSC threshold | 0.143 |
| Sharpening B-factor [Å<sup>2</sup>] | -96 |
| **Refinement** |  |
| Initial models [PDB IDs] | 6djo, 6dv1 |
| Model resolution [Å] | 3.54 |
| FSC threshold | 0.5 |
| *RMSD deviations* |  |
| Bond lengths [Å] | 0.007 |
| Bond angles [°] | 0.973 |
| Mean B-factor [Å<sup>2</sup>] | 75.1 |
| **Validation** |  |
| Molprobtity score | 2.26 |
| Clash score | 6.04 |
| *Ramachandran plot* |  |
| Favored [%] | 95.5 |
| Allowed [%] | 4.5 |
| Disallowed [%] | 0.0 |
| EMringer score | 2.71 |

**Table 2.**
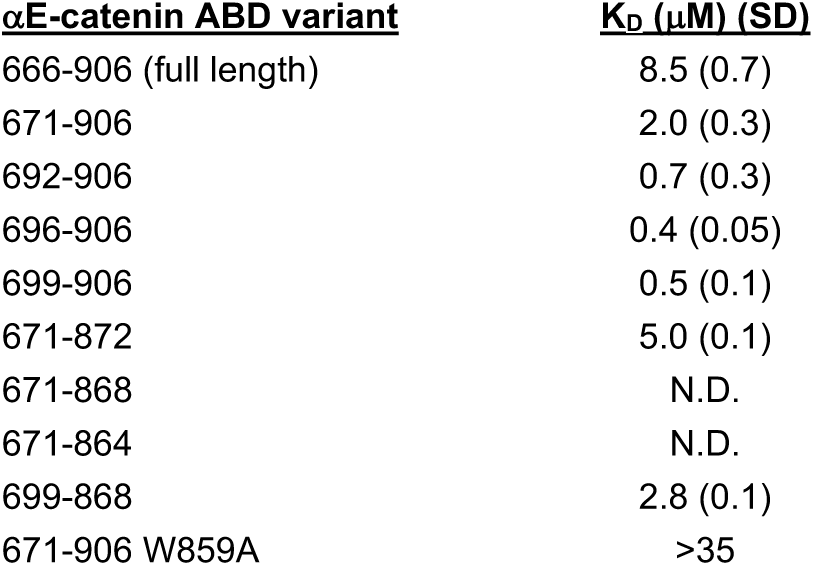
Affinities of αE-catenin ABD constructs for actin, determined by co- sedimentation. K_D_ values and standard deviations for αE-catenin 666-906, 671-906, and 692-906 are the average of 3 replicate measurements. K_D_ values for the other constructs are the average of two measurements. For αE-catenin 671-906 W859A binding did not reach saturation and therefore only a lower limit for the K_D_ is given. N.D., no detectable binding. Representative binding curves and corresponding gels are shown in Table 2-supplemental figures 1 and 2.

**Figure 2:**
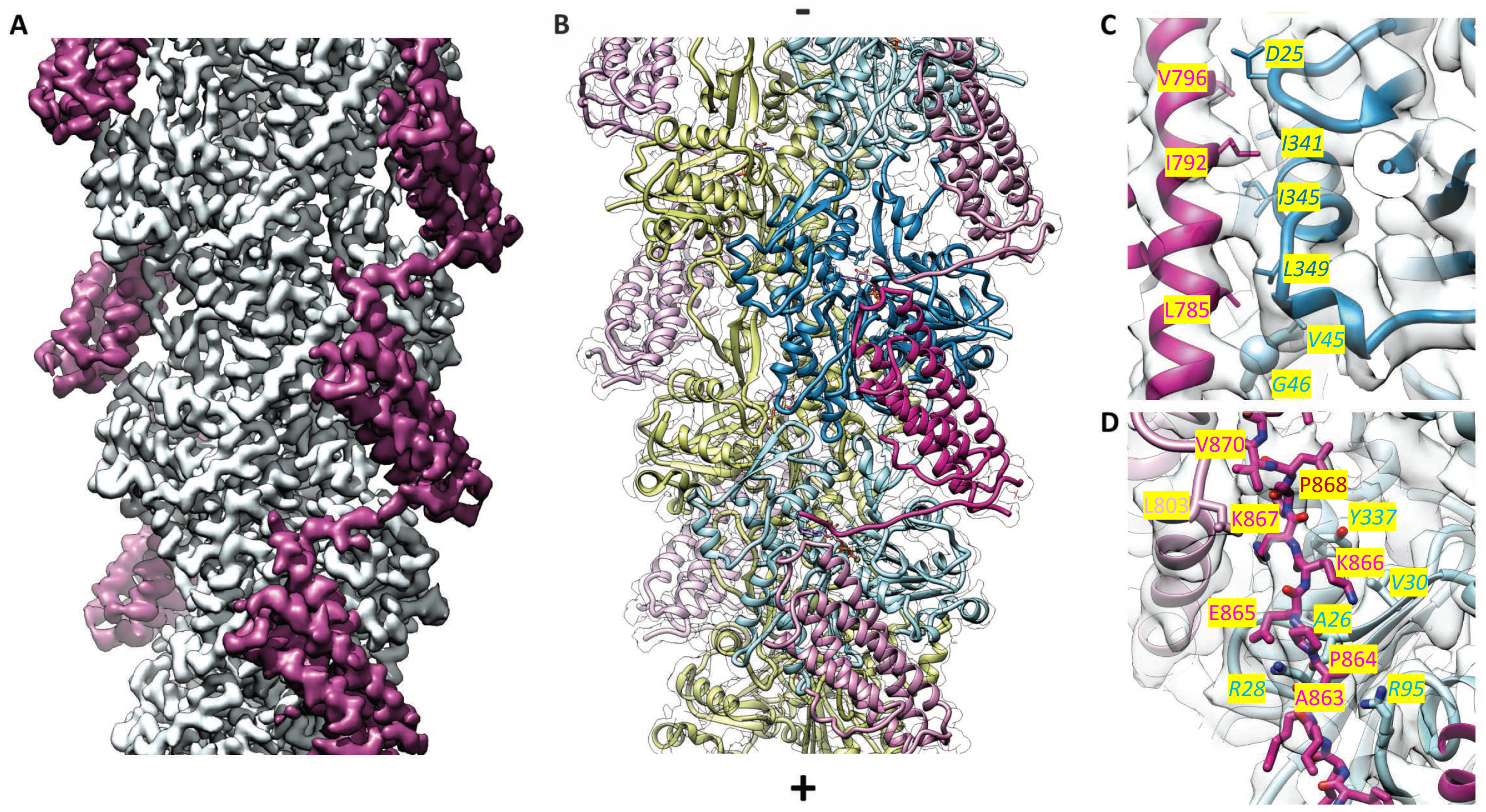
Cryo-EM analysis of the αE-catenin ABD–F-catenin complex. **A**. Cryo-EM map of the actin-ABD structure. The segmented ABDs are shown in magenta. The (-) end of the filament is shown at the top, and the (+) end at the bottom. **B**. Molecular model of a section of an actin filament bound to αE-catenin ABDs, same orientation as (A) and with transparent density map overlaid. Actin protomers are colored according to their long-pitch helix in blue and yellow. The bound ABDs are shown in magenta and pink. **C, D**. Closeups of model and cryo-EM map showing residues on H4 (panel C) and the CTE (panel D) that have been studied by site-directed mutagenesis. The ABD is shown in red, and two monomers of actin in different shades of blue. In (D), a neighboring copy of the ABD along the filament is shown in pink. Actin residue labels are *italicized*.

The cryo-EM reconstruction allowed us to build an atomic model of the complex (Figure 2), using the structures of bare ADP actin filaments (Chou and Pollard, 2019) and the αE-ABD crystal structure (Ishiyama et al., 2018) as starting points. Only the first 3 and last residues of actin could not be placed with confidence. The visible portion of the αE-catenin ABD starts at residue 699 and ends at 871; 6 residues in the loop connecting H4 and H5, and 8 residues in the connection between H5 and the rest of the C-terminal extension also could not be modeled. Local resolution analysis shows that the most well-defined region is within the actin filament core and gradually falls off towards larger radii (Fig. S1). Importantly, the interface of the ABD and F-actin is well defined in the cryo-EM map, as is the conformation of the helical bundle (Figure 2, Figure 2-figure supplement 1).

The helical rise (27.3 Å) and twist (−166.8°) of F-actin in the reconstruction are practically identical to those of bare F-actin (Chou and Pollard, 2019; Merino et al., 2018), indicating that the ABD does not induce any major changes into the filament. Consequently, the root-mean square deviation between bare ADP actin filaments and actin with the ABD bound is low (0.76 Å, Cα deviations). The most notable difference is in the conformation of the subdomain 2 loop (D-loop), a region implicated in changes associated with the ATP hydrolysis cycle (Chou and Pollard, 2019; Merino et al., 2018), stiffness and stability (Kang et al., 2012; Pospich et al., 2017), and in filament disassembly (Grintsevich et al., 2017). In the present structure, this region is in a ‘closed’ conformation similar to those observed in the ADP-bound structures (Chou and Pollard, 2019; Merino et al., 2018); modeling suggests that the ‘open’ D-loop conformation that has been associated with the ATP-bound form of bare actin filaments (Merino et al., 2018) would clash with the bound αE-catenin. Relative to the bare ADP-actin structure, however, D-loop residues 45-50 move significantly, with M47 showing the largest displacement of about 5 Å. This region contacts the αE-catenin ABD. It has been noted that tensile forces imposed on actin by the thin ice needed for cryo-EM imaging may affect the D-loop (Galkin et al., 2012), and we have previously observed similar differences at low resolution upon αE-catenin binding (Hansen et al., 2013), but whether tension has any role in the conformation observed here is unclear given the direct contacts with the ABD. Moreover, the refinement procedure used to generate high-resolution structures from cryo-EM images selects and enforces a single conformation, so it is likely that in order to achieve the highest resolution reconstruction possible, other information content including alternative conformations, was lost.

In contrast to the local changes in F-actin, the ABD undergoes large structural rearrangements upon complex formation. Compared to the unbound ABD crystal structure, the N-terminus through the last turn of H1 is disordered (Figure 3a). We note that an F-actin-bound αE-catenin ABD structure has been reported recently for the complete ABD (664-906) (Mei et al., 2020), and the same residues are disordered, demonstrating that the truncation of H0 in our construct has no influence on the actin-bound structure. The remaining four helices rearrange slightly to bind to the filament. A key part of this change involves the long loop that connects H4 and H5, where the first strand of a β-hairpin becomes a two-turn extension of H4 (residues 795-801) that forms contacts with actin (Figures 2c, 3a,b). In addition to the changes in the helical bundle, the remaining turn of H1 moves up from the bottom of the rest of the helical bundle (Figure 3c). This repositions an aromatic cluster formed by conserved residues W705 (H1), Y837 (H5), and W859 (CTE), and shifts the CTE upward (Fig. 3c). The CTE past 860, which is disordered in the isolated structure, forms an extended peptide that interacts with actin. The CTE sits between actin and the next ABD along the long pitch of the filament, and K867 directly contacts L803 of the neighboring ABD, at the top of H4 (Figure 2d). The importance of the change in the CTE is highlighted by the effect of mutating W859 to alanine, which would prevent the repacking of the aromatic cluster; this change lowers the affinity for actin approximately 10-fold (Table 1).

**Fig. 3.**
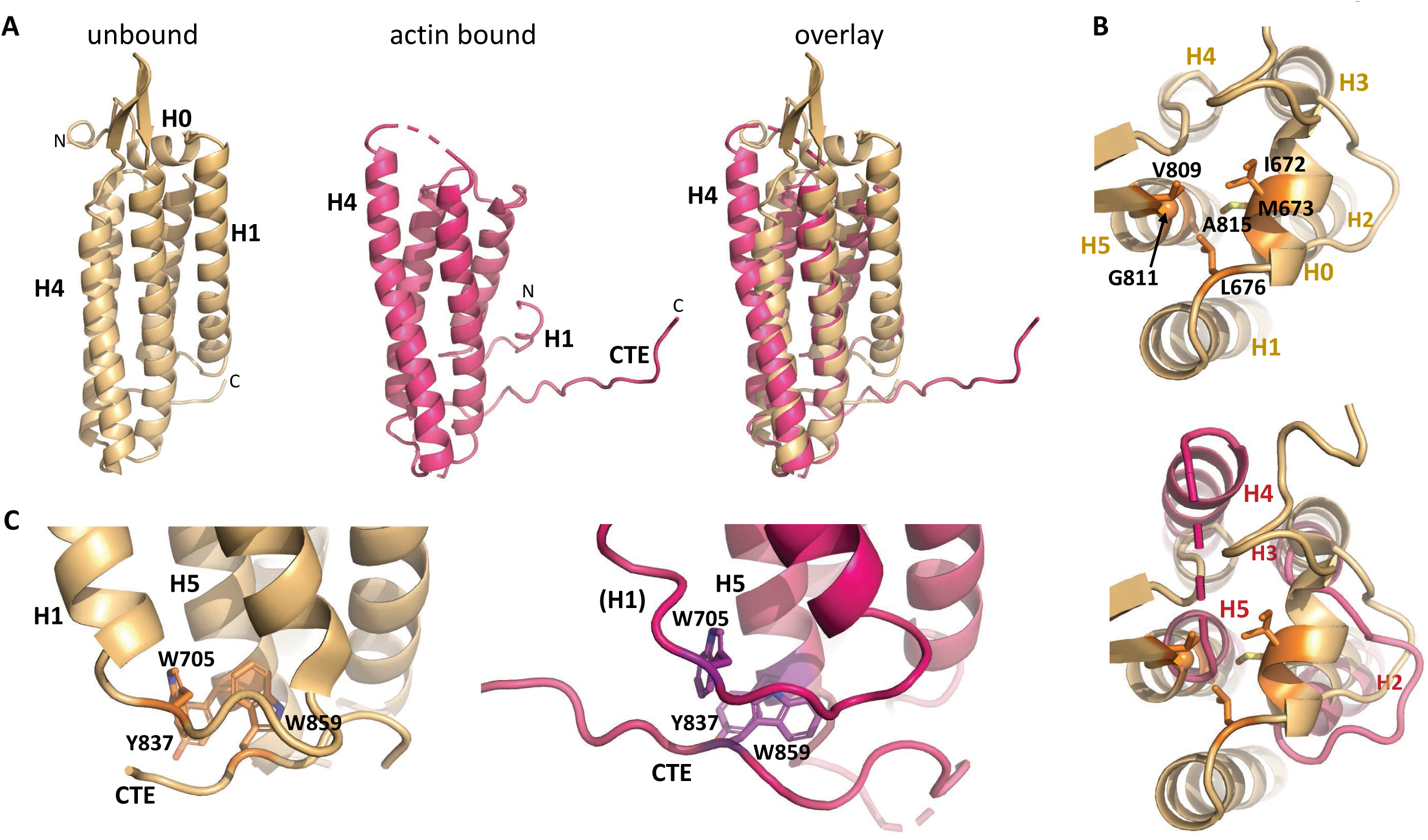
Overall changes in ABD structure upon binding to F-actin. Comparison of the unbound ABD crystal structure (orange) with the ABD in the actin-bound state (magenta). **A**. Overall comparison; the orientation is the same as Fig. 1c. **B**. Top view of H0 packing interactions lost upon its removal, and rearrangements of helices 2-5. The upper panel packing interactions of H0 residues 672, M673 and L676 with H5 residues V809, G811 and A815, and the bottom overlay shows the resulting changes in H4 and H5. **C**. Differences in the CTE. The aromatic cluster of W705 in H1, Y837 in H5, and W859 in the CTE repacks due to the shift in position of the remaining turn of H1 in the actin-bound state.

The actin-bound αE-catenin ABD complex structure is consistent with mutations that weaken its binding to F-actin. Ishiyama et al. (Ishiyama et al., 2018) mutated H4 residues L785, I792 and V796 to alanine. L785 had the most severe effect, reducing the K_D_ ∼15x; it interacts at the interface of two longitudinally adjacent actin monomers, including L349 of one monomer and V45 and G46 in the D-loop of the other (Figure 2c). I792A and V796A also had significant effects on affinity; I792 forms a packing interaction with I345 of actin and V796 packs against actin residues I341 and D25 (Figure 2c). Chen et al. (Chen et al., 2015) found several point mutants that severely weakened binding, including I792A. K842A eliminates contacts with actin residues H87 and Y91; and K866A eliminates side chain and main chain contacts with actin residues R28 and V30 (Figure 2d). Pappas and Rimm (Pappas and Rimm, 2006) removed sets of positively charged residues and saw only modest effects on binding; the largest effect was the triple mutant K747A/K748A/K797A; K797 forms a salt bridge with actin E334, whereas the other two lysine residues point into solvent on the other side of the domain.

Deletion of the C-terminal αE-catenin residues 865-906 compromises actin binding (Pokutta et al., 2002), whereas a construct ending at 883 binds (Chen et al., 2015; Pappas and Rimm, 2006). This observation is consistent with the contacts observed between actin and residues 866-868 (Figure 2d). To more precisely determine which residues of the αE-catenin CTE observed to contact actin are critical for binding, we prepared a series of C-terminal truncations of the αE-catenin 671-906 construct and compared their affinities (Table 2,Table 2-figure supplement 2). Consistent with the contacts formed by 866-868, removing residues 865-906 produced no detectable binding. Surprisingly, removal of residues 869-906 eliminated detectable F-actin binding, even though none of these residues interact with actin in our structure. Likewise, there is a slight loss of affinity upon removal of residues 873-906, even though we do not observe these residues in the structure.

The structural changes and interactions with actin observed here appear to be conserved throughout the α-catenin/vinculin family. Specifically, the residues that mediate the interactions with actin are strongly conserved throughout the α-catenin sequences (Figure 2-figure supplement 2). Moreover, although relatively few of the actin-contacting residues are conserved in vinculin (notably, those in the C-terminal portion of H4), the vinculin ABD undergoes a similar structural transition upon binding to actin (Kim et al., 2016; Mei et al., 2020).

### H0 and H1 regulate actin affinity

Despite the large changes between the free and F-actin-bound ABD structures, we see no evidence for multiple conformations of the ABD when bound to F-actin, suggesting that the 4-helix state is the stably bound one. To assess whether the rearranged state is significantly populated in solution, we compared the proteolytic sensitivity of the H0-deleted ABD used in the EM structure (671-906) in the presence and absence of F-actin, using the protease elastase. We found that in isolation the domain was resistant to digestion, whereas binding to F-actin led to the appearance of smaller, protease-resistant fragments (Figure 4a). N-terminal sequencing of the SDS-PAGE bands corresponding to these fragments revealed cleavage of H1 at residues A689 and S703 (Figure 4b). The cleavage at S703 is consistent with the very weak cryo-EM density observed between residues 699 and 702, which likely indicates that this turn of helix is flexible. The resistance of H1 to protease in the absence of F-actin implies that its association with the H2 and H5 surface is strongly favored in solution. Helix 1 of the vinculin actin-binding domain, which forms a similar four helix bundle when bound to actin (Kim et al., 2016; Mei et al., 2020), also becomes proteolytically sensitive upon binding to actin (Bakolitsa et al., 1999). The proteolysis data from both αE-catenin and vinculin suggest that the free energy of binding to actin drives the structural rearrangement of the ABD.

**Fig. 4.**
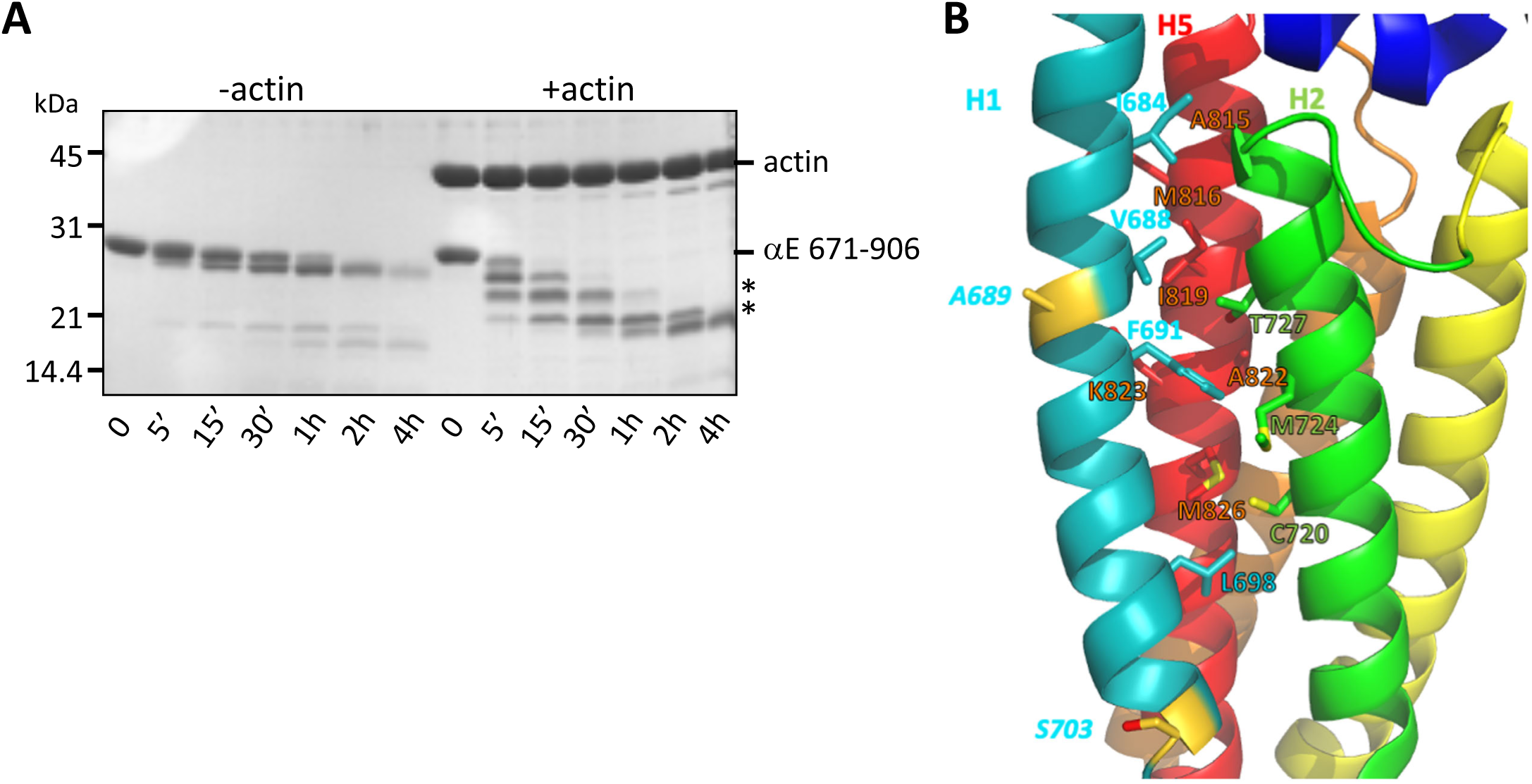
Stability of the H1- H2/H5 interface. A. Time course of elastase digestion of αE-catenin 671-906. The two smaller fragments analyzed by N-terminal sequencing are indicated with asterisks. B. Rainbow diagram of the unbound αE-catenin ABD (PDB 6dv1), colored as in Fig. 1c. The two residues at the elastase cut sites are indicated in gold. Side chains in the H1- H2/H5 interface are shown in stick representation.

If the association of H0 and H1 with the 4-helix bundle inhibits the rearrangement of the structure to a stable actin-bound conformation, they should weaken the affinity of the ABD for actin. Therefore, we prepared a series of αE-catenin ABD variants in which H0 and H1 were deleted or truncated. As more N-terminal sequence was deleted, the affinity became stronger, such that deleting H1 through residue 698 results in approximately 18x stronger binding to actin filaments relative to the complete ABD (Table 2, Table 2-figure supplement 1). Notably, Ishiyama et al. deleted H0 only and saw about a ∼3x increase in affinity (Ishiyama et al., 2018), similar to change we observed when H0 is disrupted by deleting residues 666-670 (Table 2). This observation is also consistent with the observation that the crystal structure of the αN-catenin ABD lacking H0 still forms the 5-helix assembly observed in the complete ABD (Ishiyama et al., 2018). Overall, the deletion data indicate that removal of H0 and most of H1 produce a strong actin-binding species. Notably, the enhanced binding conferred by deleting H1 can compensate for the loss of residues 869-906 (Table 2; compare 671- 868 and 699-868).

Sequence features of the ABD support the idea that the free energy of binding to actin drives dissociation of H0 and H1 and rearrangement of the remaining helices. H0 has three conserved hydrophobic residues (I672, M673 and L676) that pack against three residues at the N-terminal region of H5 (Figure 3b), two of which (V809 and G811) are poorly ordered in the actin-bound structure. H1 binds to the outer face of the H2–5 four helix bundle, interacting with a surface formed by H2 and H5. Several hydrophobic H1 residues (I684, V688, F691 and L698) are buried in this interface (Figure 4b), which would disfavor dissociation of H1. However, the H1 interaction surface formed by H2 and H5 is not strongly hydrophobic, comprising four methionine residues (M723, M724, M816 and M826), C720, T727, I819, A822 and the aliphatic portion of K823 (Figure 4b; Figure 2-Figure supplement 2), which suggests that there would not be a large destabilization of the 4-helix bundle upon removing H1 from this surface. Indeed, the construct starting at 698, which deletes all of the H1 sequence missing in the EM structure, is well behaved in solution (it is monomeric as assayed by size exclusion chromatography-coupled multi-angle light scattering; data not shown), consistent with the idea that exposure of this surface is not energetically disfavored. Notably, the mildly hydrophobic character of this H2/H5 surface is strongly conserved throughout the α-catenin family (Figure 2-Figure supplement 2).

### Insights into catch bond mechanism

Our structural and biochemical data indicate that removal of H0 and H1 enable the structural transition of the C-terminal half of H4 and movement of the CTE (Figure 3), which result in additional contacts with F-actin and stable binding (Figure 4). Given that the vinculin ABD lacks H0 but its structure bound to actin shows the same four-helix, rearranged bundle relative to vinulin in solution (Mei et al., 2020), and that the crystal structure of the αN-catenin ABD lacking H0 retains the five helix bundle architecture of the full ABD (Ishiyama et al., 2018), it is clear that removal of H1 is the major determinant in achieving the high affinity actin-binding state. The binding data in solution, *i*.*e*., in the absence of applied force, demonstrate that the free energy of binding to actin drives this transition. Optical trap data from both αE-catenin and vinculin indicate that force enhances the stability (specifically, the bound lifetime) of the ABD-actin interaction. Modeling of the optical trap data indicated that force prevents the transition from a strongly bound to a weakly bound state, as well as promoting the transition to a strongly bound state (Buckley et al., 2014; Huang et al., 2017). Thus, while force is not needed to stably bind actin, it enhances the strength of the ABD-actin interaction by shifting the equilibrium between weakly and strongly bound states toward the strong state.

To gain structural insight into how force affects the weak and strong actin-binding states of the ABD, we superimposed the isolated αE-catenin ABD structure on the actin-bound version (Figure 5a). This revealed only minor clashes with actin at the N-terminus of H5, and modeling suggests these can be alleviated by changes of side chain rotamers (Figure 5a, left). Moreover, as the N-terminal half of H4 and almost all of H5 superimpose closely (Figure 5a, middle), it is likely that in the five-helix conformation these regions could form interactions similar to those visualized in the EM structure. However, key interactions made by the C-terminal part of H4, including those of I792 and V796 (Figure 2c), would not form. These observations may indicate that the 5-helix conformation can bind F-actin weakly, and we propose that this conformation represents the weak binding state observed in the optical trap (Figure 5b). We cannot assess whether the weak state corresponds precisely to the solution conformation, or if it is intermediate between the experimentally determined unbound and actin-bound structures.

**Fig. 5.**
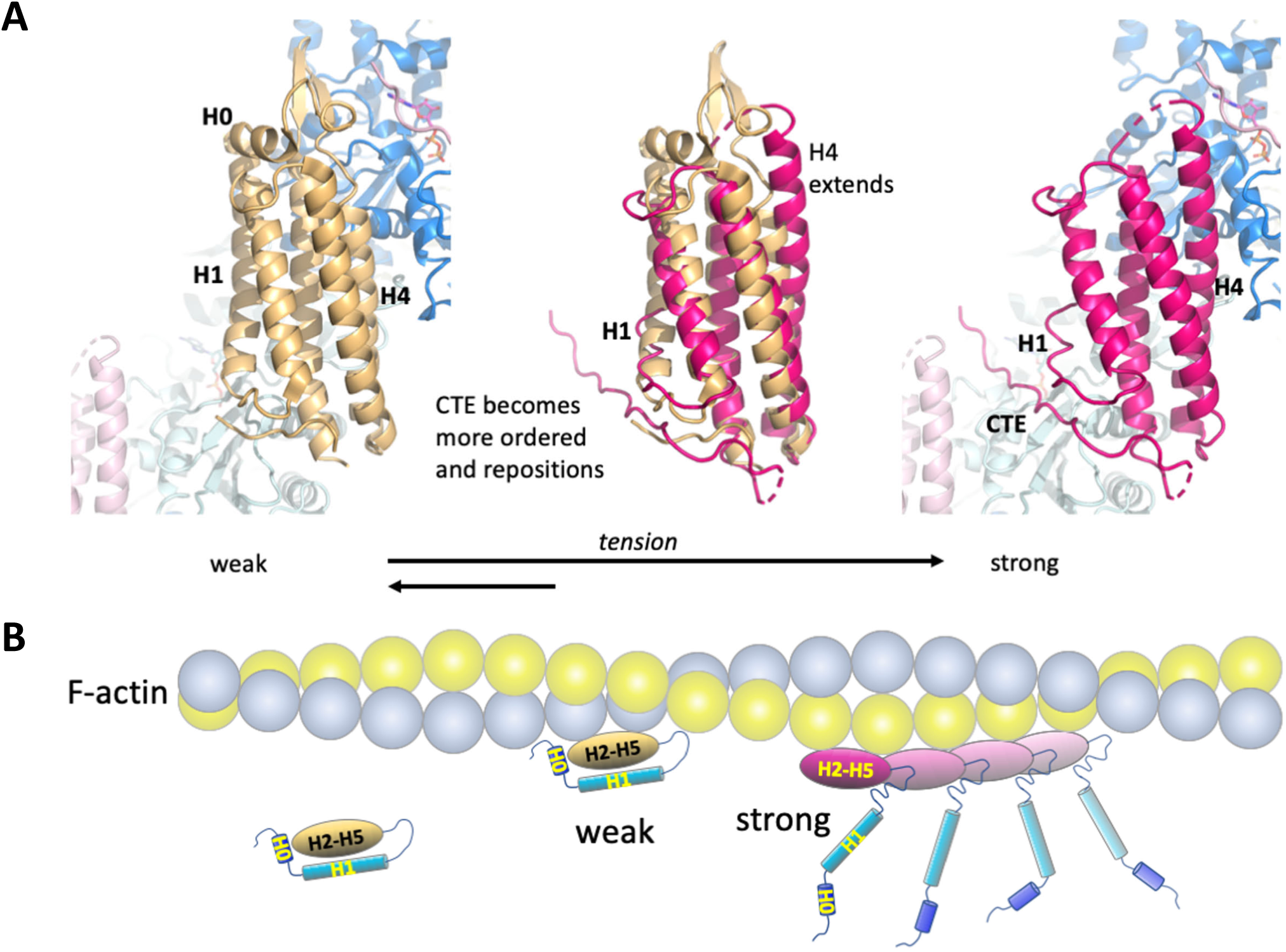
Model of the weak and strong actin-binding states of αE-catenin. A. Superposition of the isolated αE-catenin ABD on the actin-bound structure reveals no major clashes with F-actin (left panel). When the ABD is bound to F-actin, H0 and H1 dissociate from the H2-5 bundle, which results in the extension and shift of the C-terminal part of H4 as well as ordering and repositioning of the CTE to bind to actin. B. Schematic diagram of αE-catenin ABD conformational states when unbound, weakly bound, and strongly bound to actin. Cooperative binding of the ABD, as observed in the cryo-EM structure, is illustrated for the strong state.

Assuming that the 4-helix conformation observed in the complex with F-actin represents the strong state, it is likely that force on the ABD prevents H1 and H0 from associating with the rest of the ABD (Figure 6a). Tension on the N-terminus of the ABD that is stably bound to F-actin will prevent re-association of H1 and H0, thereby favoring the strongly bound state and enhancing its lifetime. Conversely, if the 5-helix conformation binds weakly, force would provide additional energy to drive the dissociation of H0 and H1 from the bundle (Figure 6b), thereby facilitating the transition to the strongly bound state. We also note that the cooperative binding of the ABD to F-actin, which may be due to the ordering of the CTE in the 4-helix conformation and its interaction with the longitudinal ABD neighbor on actin (Figure 2d), likely has a role in the stable binding (Figure 5b).

**Fig. 6.**
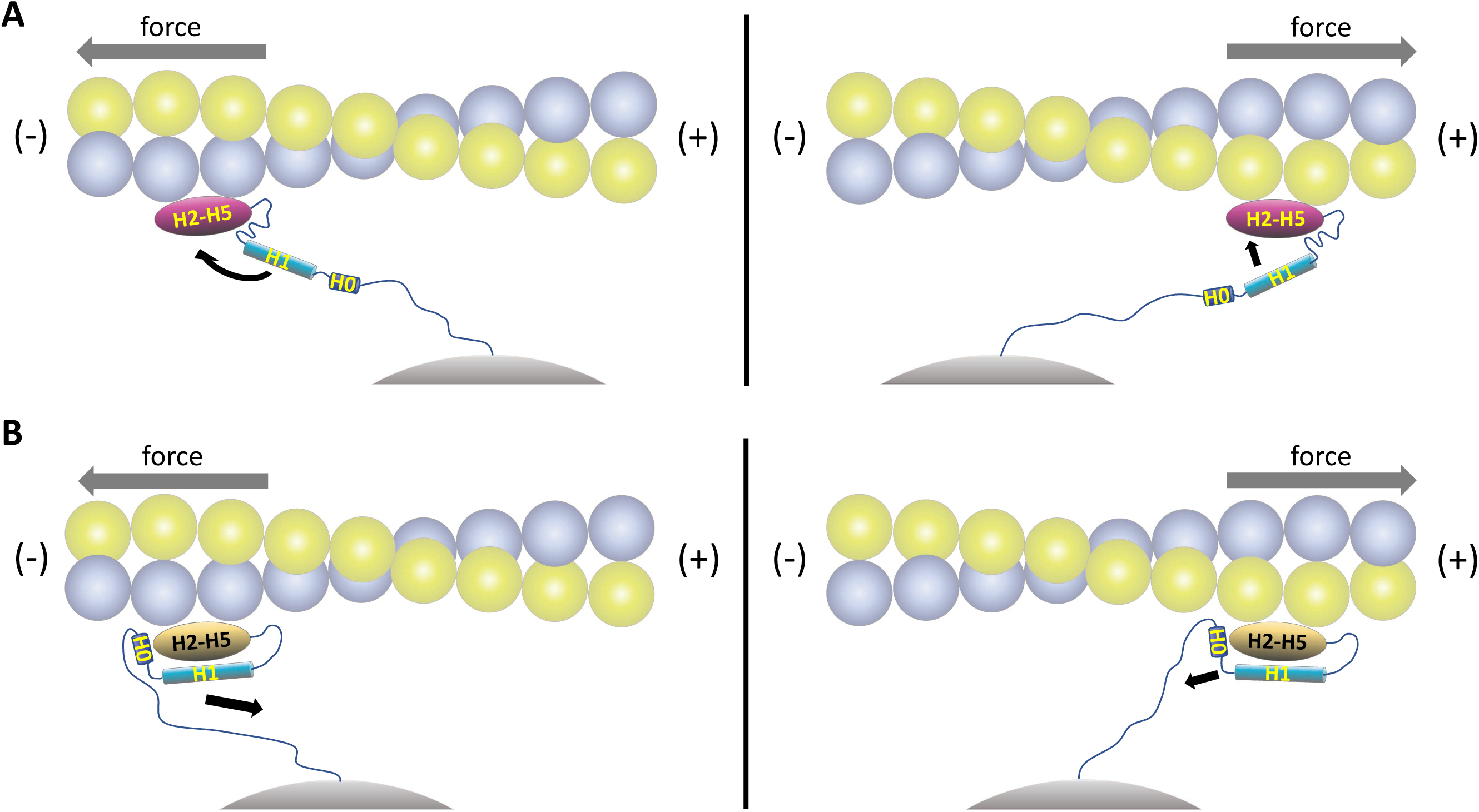
Model of directional catch bonding. Actin and the αE-catenin ABD are illustrated as in Fig. 5b. The N-terminus of the ABD is shown tethered to a stationary point, *i*.*e*., as part of the cadherin/β-catenin/α-catenin complex. The grey arrows indicate the direction of force. A. Tension applied to the bound strong state prevents re-binding of H0/H1 to the H2-H5 bundle. Force applied in the (-) direction will move H1 away from the H2-H5 bundle and place H1 in an unfavorable orientation for rebinding, whereas force directed in the (+) direction will place H1 closer to and in a more favorable orientation for rebinding. B. Tension applied to the bound weak state will remove H0/H1 from the H2-H5 bundle. Force applied in the (-) direction will tend to pull H0/H1 away from the H2-H5 bundle, whereas force in the (+) direction is predicted to have a smaller effect on H0/H1 dissociation.

## DISCUSSION

Catch bond behavior has been observed in a variety of proteins subject to mechanical force. Examples include the extracellular portions of cell adhesion molecules such as bacterial FimH that attaches to the urinary tract epithelium, selectins that mediate rolling of leukocytes on endothelia, integrins that mediate cell-extracellular matrix adhesion, and classical cadherins (Pruitt et al., 2014). For FimH, selectins and integrins, hinges between domains change position under mechanical load, and these changes can be transmitted to binding domains in a variety of ways to promote a strong ligand-binding state (Le Trong et al., 2010; Pruitt et al., 2014; Rakshit et al., 2012; Springer, 2009; Springer et al., 2008; Xiao et al., 2004). More recent work has revealed catch bonding to F-actin by the intracellular adhesion proteins αE-catenin and vinculin. These proteins also have multiple domains that are likely to change their relative positions upon application of force, as has been demonstrated for αE-catenin (Barrick et al., 2018; Choi et al., 2012; Kim et al., 2016; Li et al., 2015; Terekhova et al., 2019). Attachment of the α-catenin N-terminal domain to β-catenin and the C-terminal ABD to actin (Figure 1b) implies that tension is transmitted through the entire protein.

The structure of the actin-bound αE-catenin ABD presented here provides a molecular-level explanation of the catch bond behavior of αE-catenin as well as the homologous vinculin (Abore et al., 2020; Buckley et al., 2014; Huang et al., 2017). By serving as a bridge between β-catenin and actin, αE-catenin is placed under tension, which likely stretches the linker between the M domain and the ABD and thereby applies tension to H0 and H1. Likewise, binding of vinculin to talin and actin in focal adhesions will stretch the loop that precedes the ABD. Tension stabilizes the 4-helix, strong-binding conformation of these ABDs bound to F-actin by preventing rebinding of H1 (and H0 in the case of αE-catenin). The stabilization of the αE-catenin CTE in the 4-helix conformation is likely an important component of the strongly bound state: it not only forms interactions with actin but also contacts a neighboring ABD on the filament, which may underlie its cooperative binding to F-actin (Buckley et al., 2014; Hansen et al., 2013). Moreover, modeling of the isolated αE-catenin ABD structure on actin shows few clashes but suggests that the 5-helix conformation forms a subset of the interactions observed in the cryo-EM structure, and this likely corresponds to the weakly bound state. If so, then tension applied to the weakly bound state could facilitate dissociation of H0 and H1 and therefore transition to the strongly bound conformation (Figure 6).

Optical trap and biolayer interferometry measurements indicate that in the absence of force, 80-90% of the actin-bound αE-catenin molecules are in the weak state (Buckley et al., 2014; Ishiyama et al., 2018). The presence of a small fraction in the strong state implies that the free energy of binding to actin promotes dissociation of H1 even in the absence of force (Figure 6a). Our deletion data indicate that removal of H1 does not destabilize the rest of the ABD, but the burial of several hydrophobic H1 residues in the H2/H5 interface (Figure 4b) suggests that H1 dissociation would be disfavored. The energy input by mechanical force acting over a certain distance helps to overcome this barrier.

A recent study proposed that removing H0 enables the strongly bound, force-enhanced state of αE-catenin (Ishiyama et al., 2018), although the structure of the actin-bound ABD showing the dissociated H1 and rearranged H2-H5 region was not available to these authors. Their proposal was based in part on the observation that in order to fit biolayer interferometry F-actin binding data for the complete ABD, a model invoking two species with different affinities was needed (Ishiyama et al., 2018). The increased affinity without H0 ((Ishiyama et al., 2018); Table 2) does show that the stabilization energy provided by the interaction of H0 with the rest of the molecule contributes to the barrier to the transition to the stably bound conformation in αE-catenin, as H0 likely interferes with the rearrangements of H2–H5 (Fig. 3b). However, the fact that the crystal structure of a mutant αN-catenin ABD lacking H0 is not significantly different from the crystal structure of the complete ABD (Ishiyama et al., 2018) indicates that removing only H0 does not produce the rearranged, strong-binding 4-helix bundle conformation in solution. Moreover, the vinculin ABD lacks H0, yet also forms catch bonds with actin and forms a 4-helix bundle similar to that of αE-catenin when bound to F-actin (Huang et al., 2017; Kim et al., 2016; Mei et al., 2020). Also, both the vinculin ABD and αE-catenin ABD lacking H0 are resistant to protease digestion when not bound to actin (Figure 4; (Bakolitsa et al., 1999)). Finally, removal of H0 and H1 produces a significantly higher affinity for F-actin than removing only H0 (Table 2). These observations imply that dissociation of H1 is the major factor that determines stable binding to actin. The free energy of binding to F-actin drives H1 dissociation, and mechanical force further enhances the dissociated state.

The catch bonds formed by αE-catenin, vinculin and their ABDs with actin show a strong asymmetry (Abore et al., 2020; Huang et al., 2017) (N. Bax, D. Huang, A. Wang, A. Dunn, and W.I.W., manuscript in preparation). Specifically, force directed toward the (-) end of the filament greatly enhances the strongly bound, long-lived state, whereas force towards the (+) has a more modest effect on bound lifetimes. (We originally reported (Buckley et al., 2014) that the catch bonding of the cadherin/β-catenin/αE-catenin complex was not asymmetric, but this proved to be due to limitations in the sensitivity of the instrument used in that study; N. Bax, D. Huang, A. Wang, A. Dunn, and W.I.W., manuscript in preparation). The present structure offers a clue to this behavior (Figure 6). With the rest of α-catenin or vinculin bound to a stationary anchor (*i*.*e*., cadherin/β-catenin or integrin/talin), force in the (-) direction of the filament would result in pulling the N-terminus of the ABD in the opposite direction, positioning H1 away from the H2-5 bundle and disfavoring its rebinding (Figure 6a). Likewise, force applied to the weakly-bound 5 helix conformation would result in H1 “peeling off” from the bundle. Force applied in the opposite (+) direction would tend to move the N-terminus of the ABD such that H1 would be more aligned with its orientation found in the 5-helix state, giving it a higher probability of rebinding to the H2-5 bundle (Figure 6b). Thus, a smaller effect of force in stabilizing the strong state would be observed in this direction. While myosin II contractility also applies force in the (-) direction of actin filaments, the actual direction of the applied force vectors for both αE-catenin and vinculin depends on the detailed geometry of the full adhesive complexes and their organization in the cell. Determining these parameters will be needed to fully understand catch bonding by these proteins.

## METHODS

### Expression and purification of αE-catenin ABD constructs

αE-catenin ABD constructs were cloned into a pGEX-4T-3 or pGEX-TEV bacterial expression vector; the latter is a modified pGEX-KG vector with a tobacco etch virus (TEV) protease recognition site inserted after the GST-tag and the thrombin cleavage site. N-terminal GST-fusion proteins were expressed in *E. coli* BL21cells. Cells were grown at 37°C to an OD_600_ of 0.8-1.0 and induced overnight at 18°C with 0.5 mM isopropyl-1-thio-β-d-galactopyranoside. Cells were harvested by centrifugation and pellets were resuspended in 20 mM Tris pH 8.0, 200 mM NaCl and 1mM DTT. Before lysis in an Emusliflex (Avastin), protease inhibitor cocktail (Mixture Set V, Calbiochem) and DNase (Sigma) were added. After centrifugation at 38,500 × *g* for 30 min the lysate was incubated with glutathione-agarose beads for 1 h at 4°C. After washing the beads with PBS containing 1 M NaCl and 1 mM DTT, the beads were equilibrated with either 20 mM Tris pH 8.5, 150 mM NaCl, 1 mM DTT for thrombin cleavage or 20 mM Tris pH 8.0, 150 mM NaCl, 1 mM DTT, 1 mM EDTA, 10% glycerol for TEV cleavage. The protein was cleaved for 2h at room temp with thrombin or overnight at 4°C with TEV. Cleaved protein was eluted from the beads and further purified on a cation exchange column (Mono S 10/100, GE Healthcare) in MES pH 6.5, 1mM DTT buffer with a 0-500 mM NaCl gradient, followed by size exclusion chromatography (Superdex S200, GE Healthcare) in 20 mM HEPES pH 8.0, 150 mM NaCl, 1mM DTT.

### Actin binding assay

G-actin prepared from rabbit muscle (Spudich and Watt, 1971) was stored in 40 μM aliquots at -80°C. Frozen aliquots were thawed on ice and centrifuged for 20 min at 140717 x *g* in a Beckman TLA 100 rotor. After centrifugation the concentration was determined by UV absorbance at 290 nm and G-actin was polymerized by addition of 10x F-buffer (100 mM pH 7.5 Tris, 500 mM KCl, 20 mM MgCl2, 10 mM ATP) and incubation for 1 h at room temperature. Aliquots from the same batch of actin were used for all polymerization assays, and efficient polymerization was confirmed by pelleting at 20 min at 140,717 x *g* and analysis of the supernatant and pellet by SDS page. F-actin was stored for up to two weeks at 4°C. For sedimentation assays, F-actin was diluted to 4 μM with buffer A (20 mM HEPES pH 8.0, 150 mM NaCl, 1mM DTT, 2mM MgCl_2_, 0.5 mM ATP, 1mM EDTA). A dilution series of purified αE-catenin ABD in 20 mM HEPES pH 8.0, 150 mM NaCl and 1 mM DTT was set up and an equal volume of 4 μM F-actin or buffer A was added. The mixture was incubated for 30 min at room temperature. Samples were centrifuged in a Beckman TLA100 rotor at 140717 x *g* for 20 min at 4°C. The supernatant was carefully removed and the pellet resuspended in reducing Laemmli buffer. Samples were run on SDS PAGE. Coomassie-stained bands were scanned and quantified on a LI-COR Odyssey scanner (LI-COR Biosciences). To extrapolate concentration from band intensity a dilution series of αE-catenin ABD was run in parallel for each assay and stained and destained under the same conditions as the assay itself. To correct for SDS-PAGE loading errors, for each concentration of αE-catenin, its band intensity was normalized by the ratio of the actin band intensity at that point and the average actin band intensity calculated over all concentration points. The data were analyzed in the program GraphPad Prism (GraphPad Software, La Jolla California USA) and fitted with a ‘single binding with Hill coefficient’ model. The data were analyzed in the program GraphPad Prism and fitted using the ‘specific binding with Hill slope’ model, with the exception of the αE-catenin 671-906 W859A mutant. In that case, the curves did not reach saturation, and fitting with a Hill coefficient was not possible, so a ‘One site-specific binding’ model was used to obtain K_D_ estimates. In this case the binding is sufficiently weak that we report a lower limit on the K_D_ rather than a specific value (Table 2).

### Electron cryo-microscopy sample preparation

Rabbit skeletal actin was prepared as described (Kang et al., 2012; Spudich and Watt, 1971) and was used within 1 week of preparation. Fresh complete (residues 666- 906) or truncated αE-catenin-ABD (residues 671-906) were used within 1-2 days of preparation. Both filamentous actin and the respective αE-catenin ABD construct were diluted into KMEI buffer (10 mM Imidazole pH 7, 50 mM KCI 2 mM MgCI_2_, 1mM EGTA, 0.2 mM ATP, 2 mM DTT) at 0.125 mg/ml actin and 0.25 mg/ml ABD, corresponding to an ABD concentration of 10 μM. After 10 minutes of incubation, 5 or 4 μl from the final 1:2 (wt/wt) mixture was applied to plasma cleaned C-flat copper grids 2/1 or 2/2 (Protochips Inc.), respectively. After 1 minute of incubation in a humidified chamber, excess liquid was manually blotted, and the samples were plunge-frozen in liquid nitrogen-cooled liquefied ethane using an in-house designed cryo-plunger.

Screening for the best sample mixture ratios and blotting conditions was performed on a T12 Tecnai Spirit electron microscope (ThermoFisher Scientific) equipped with a 4Kx4K Eagle camera (ThermoFisher Scientific), operated at a voltage of 120 kV. Data sets were acquired on Titan Krios electron microscope (ThermoFisher Scientific) equipped with an XFEG and operated at a voltage of 300 kV. Although the sample preparation protocol was optimized, we had to screen for usable grids and grid squares manually. Images were recorded with a back-thinned 4K×4K Falcon 2 (ThermoFisher Scientific) direct detection camera under minimal dose conditions using the automatic data collection software EPU (ThermoFisher Scientific). Within each selected grid hole, two positions were imaged, each with a total exposure of 1 s. A total of 5573 dose-fractionated image stacks with seven frames each were collected with a 1.035-Å pixel size at defoci ranging from -0.8 µm to -2.8 µm.

### Cryo-EM image processing

Dose weighting and motion correction were applied using MotionCor2 (Li et al., 2013) with 5 × 5 patches. The defocus was estimated either using Gctf (Zhang, 2016) or CTFFIND4 (Rohou and Grigorieff, 2015). 804 images were discarded during real-time screening at data collection time for excessive drift, strong astigmatism, and low visibility of Thon rings. The remainder was processed with the helical reconstruction routines in RELION3 (He and Scheres, 2017; Scheres, 2012; Zivanov et al., 2018). For the truncated αE-catenin ABD (671-906) bound to rabbit actin, a total of 728,331 filament segments were extracted using a box size of 200 × 200 pixels from 63,480 manually picked filaments. Two-dimensional reference-free classification for the data set was carried out in RELION3 to eliminate bad segments, reducing the number of segments from 728,331 to 422,822. An in-house rabbit skeletal actin filament reconstruction filtered to 40 Å resolution was used for an initial model. After several rounds of refinement and manual removal of bad particles and enrichment of segments showing clear decoration, the estimated resolution of the reconstruction, using the 0.143 FSC cutoff gold-standard procedure implemented in RELION3, reached 3.56 Å after postprocessing. The helical rise of the reconstruction was 27.3 Å with a helical twist of - 166.77°. The postprocessing consisted of applying the RELION3-based B-factor sharpening and application of a soft-edged mask generated in RELION3 corrected for helical edge effects using pyCoAn, an extended python version of CoAn (Volkmann and Hanein, 1999). Additional sharpening was applied using pyCoAn. The reconstruction was then symmetrized within pyCoAn using the refined helical parameters. Real space refinement was performed with Phenix (Afonine et al., 2018). Local resolution estimates were calculated with ResMap (Kucukelbir et al., 2014) and RELION3.

A molecular model was produced starting from a structure of bare ADP actin filaments (PDB code 6djo) and the αE-ABD crystal structure (PDB code 6dv1), then iteratively adjusted manually with Coot (Emsley et al., 2010) and subjected to real-space refinement in Phenix (Afonine et al., 2018). Quality indicators, including MolProbity (Chen et al., 2010) and EmRinger (Barad et al., 2015) scores, were calculated with Phenix.

The coordinates and cryo-EM map of the αE-catenin–F-actin complex have been deposited in the Protein Data Bank, identifiers 6WVT and EM EMD-21925, respectively.

### Limited proteolysis

Limited proteolysis of αE-catenin 671-906 was performed with elastase (Worthington Biochemical). αE-catenin ABD (8 μM) was incubated in the presence or absence of 8 μM F-actin for 1 h at room temperature in 5mM Tris pH 8.0, 50 mM potassium chloride, 2 mM magnesium chloride and 0.5 mM DTT. After addition of elastase to a final concentration of 0.009 mg/ml, aliquots were removed after 5’, 15’, 30’, 1h, 2h, 4h and the proteolysis reaction was stopped by addition of SDS sample buffer and boiling. Samples were run on SDS PAGE and gels were stained with Coomassie Blue. For N-terminal sequencing, bands were transferred on a PVDF membrane. Bands of ∼24 and ∼26 kDa found only in the presence of F-actin were excised and submitted for N-terminal (Edman) protein sequencing.

## Acknowledgements

We thank Max Pokutta for technical assistance. W.I.W. thanks Alex Dunn for discussions. This work was supported by U.S. National Institutes of Health grant GM118326 (D.H., N.V. and W.I.W.) and GM131747 (W.I.W.). NIH grants S10-OD012372 and S10-OD026926, and PEW innovation funds 864K625 to D.H. funded the purchase of the Titan Krios electron cryo-microscope (FEI/Thermo Fisher), Falcon II and Falcon III direct detector (FEI/Thermo Fisher), and upgrades for both the T12 and Titan Krios hardware and software including the Eagle CMOS imaging device on the T12.

## SUPPLEMENTAL FIGURE LEGENDS

**Figure 2-figure supplement 1.**
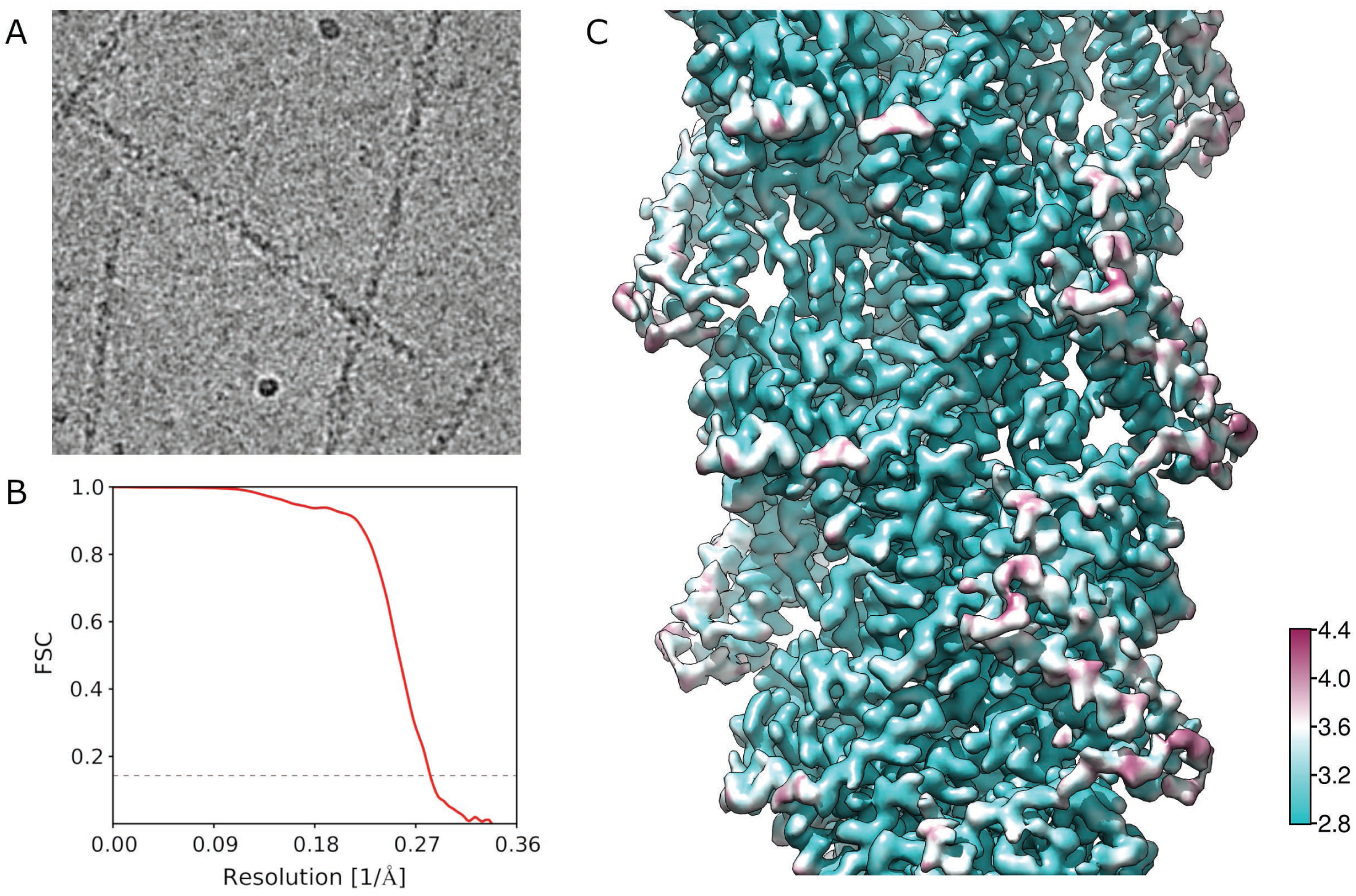
Cryo EM analysis. A. Representative image of ABD-bound filaments. B. Fourier Shell Correlation (FSC) curves using two reconstructions independently derived from two halves of the data. The 0.143 FSC cutoff used for estimating the resolution is indicated as a dashed line. C. Local resolution analysis.

**Figure 2-figure supplement 2.**
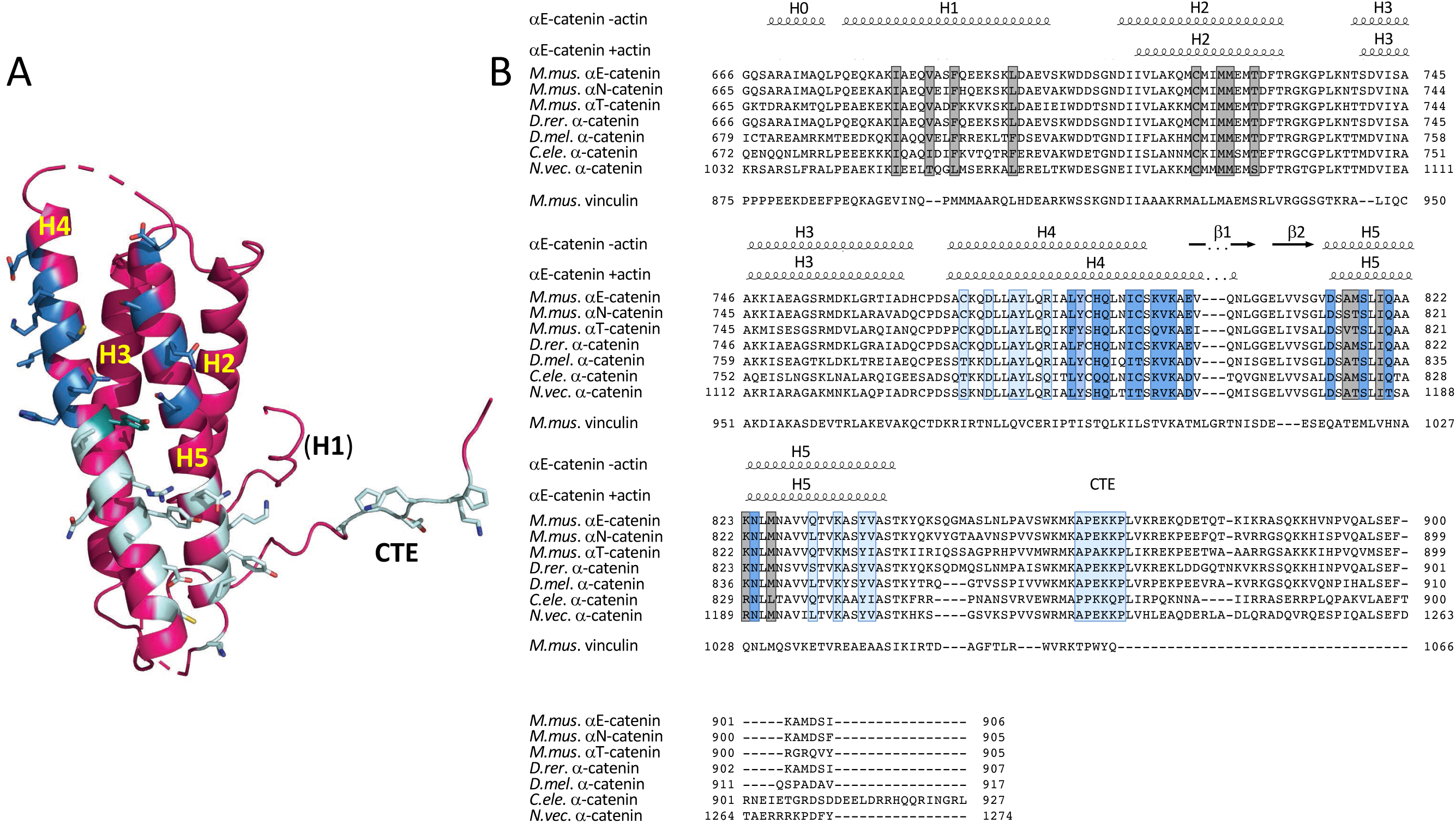
Alignment of α-catenin ABD sequences. A. Residues that form contacts with the two actin protomers (colored as in Fig. 2) are shown in stick representation. B. ABD sequence alignments. The secondary structure elements of the unbound ABD and the actin-bound ABD are shown above the alignments. Residues that contact actin are highlighted on darker and lighter blue according to which actin protomer they contact as shown in panel A and Fig. 2. Residues in grey form the interface between H1 and H2/H5. The sequence alignment was done in Geneious 10.2.2 (https://www.geneious.com). The figure was prepared with ENDscript (Robert and Gouet, 2014) and UCSF Chimera (Pettersen et al., 2004). Abbreviations used *M*.*mus*.*-Mus musculus, D*.*rer. - Danio rerio, D*.*mel. – Drosopila melanogaster, C. ele*.*-Caenorhabditis elegans, N*.*vec. - Nematostella vectensis*.

**Table 2-figure supplement 1.**
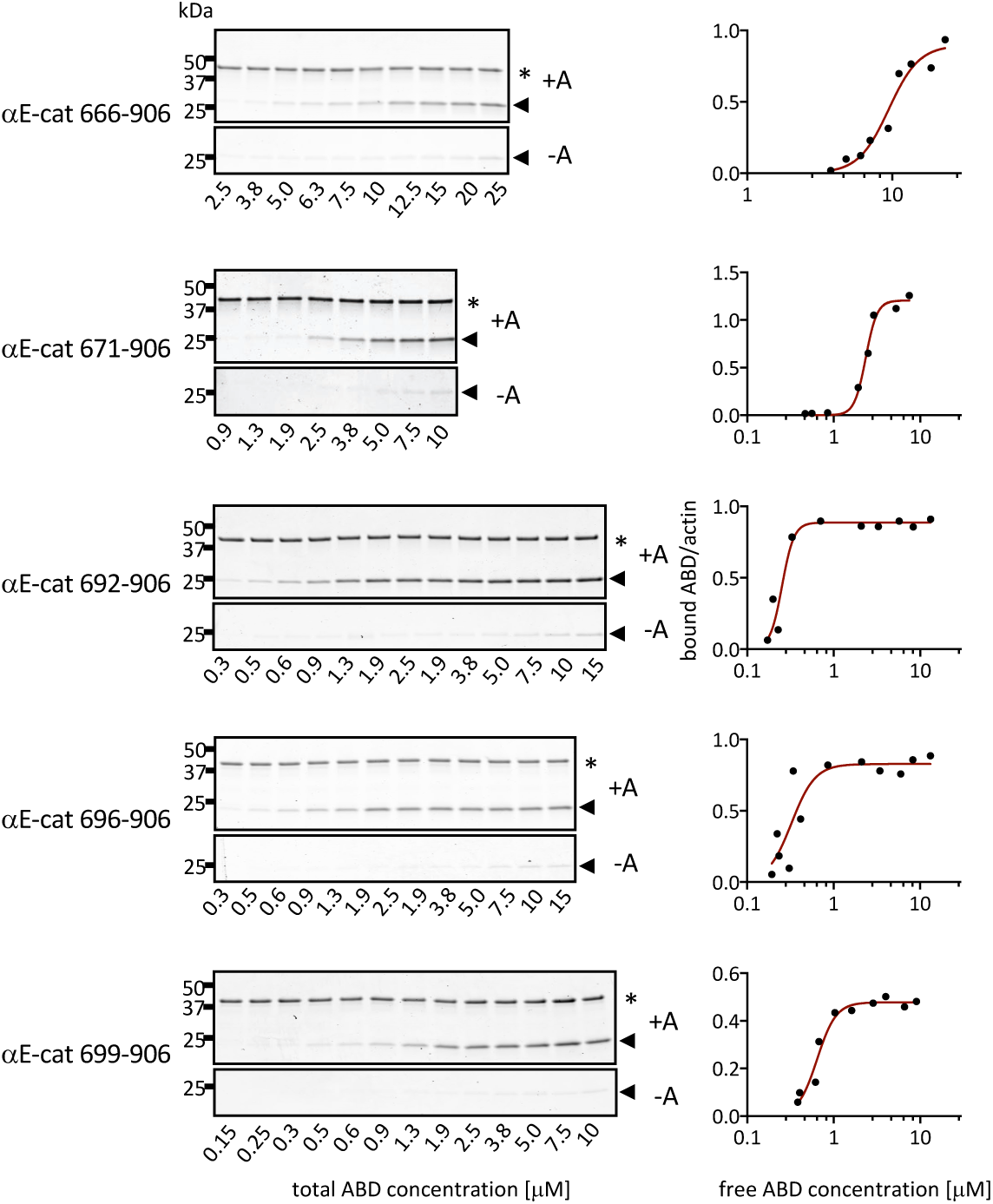
Representative gels and binding curves for actin co-sedimentation assays with αE-catenin C-terminal deletion constructs. Pellets of actin co-sedimentation assays with (+A) and without (-A) F-actin are shown. * indicates the actin band and the arrow indicates the ABD band.

**Table 2-figure supplement 2.**
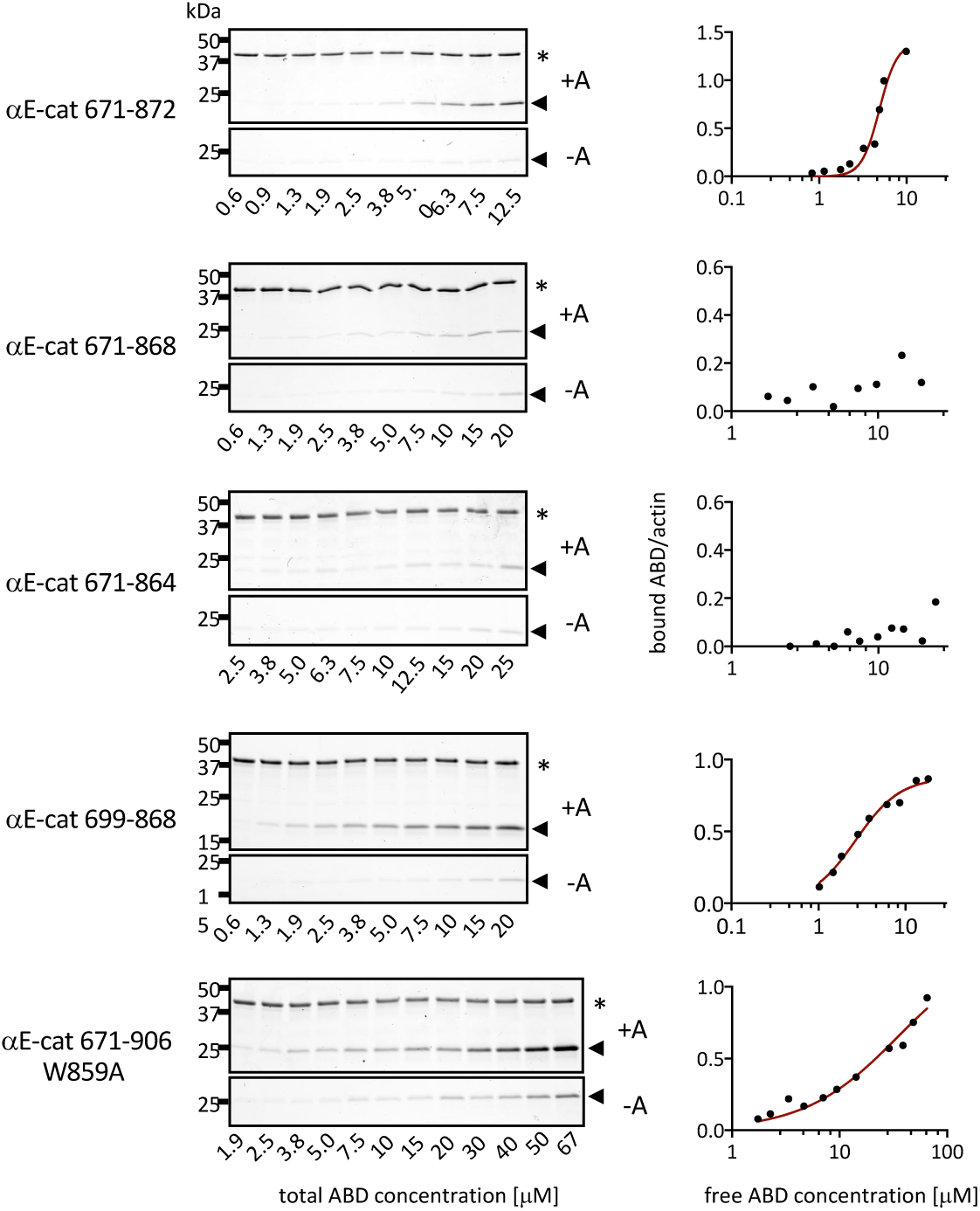
Representative gels and binding curves for actin co-sedimentation assays with αE-catenin N-terminal deletion constructs. Pellets of actin co-sedimentation assays with (+A) and without (-A) F-actin are shown. * indicates the actin band and the arrow indicates the ABD band.

